# Toxic eburicol accumulation drives the antifungal activity of azoles against *Aspergillus fumigatus*

**DOI:** 10.1101/2024.03.02.582832

**Authors:** Hesham Elsaman, Evgeny Golubtsov, Sean Brazil, Natanya Ng, Isabel Klugherz, Karl Dichtl, Christoph Müller, Johannes Wagener

## Abstract

Azole antifungals inhibit the sterol C14-demethylase (CYP51/Erg11), a key enzyme in the ergosterol biosynthesis pathway. They have fungistatic effects against yeasts but fungicidal effects against molds. The molecular basis for this difference remained unknown. The sequence of enzymatic steps required for ergosterol biosynthesis is different in yeasts and molds. Here we show that the azole-induced synthesis of fungicidal cell wall carbohydrate patches in the pathogenic mold *Aspergillus fumigatus* strictly correlates with the accumulation of the CYP51 substrate eburicol. A lack of other essential ergosterol biosynthesis enzymes, such as sterol C24-methyltransferase (Erg6A), squalene synthase (Erg9) or squalene epoxidase (Erg1) does not result in comparable cell wall alterations. Partial repression of Erg6A, which converts lanosterol into eburicol, increases azole resistance. The sterol C5-desaturase (ERG3)-dependent conversion of eburicol into 14-methylergosta-8,24(28)-dien-3β,6α-diol, the “toxic diol” responsible for the antifungal effects of azoles in yeasts, is not required for the fungicidal effects in *A. fumigatus*. In contrast to yeast, where a lack of ERG3 functionality causes azole resistance, *A. fumigatus* lacking ERG3 becomes more azole susceptible. Mutants lacking mitochondrial complex III functionality, which are less susceptible to the fungicidal effects of azoles, but get strongly inhibited in growth, convert eburicol much more efficiently into the supposedly “toxic diol”. Our results support a mechanistic model where the mode of action of azoles against the pathogenic mold *A. fumigatus*, other than in yeast, relies on the accumulation of eburicol which exerts fungicidal effects by triggering the formation of cell wall carbohydrate patches.

## INTRODUCTION

Azole antifungals are widely used as biocides, plant protection products and antimicrobials for chemoprevention, chemoprophylaxis and treatment of fungal infections in animals and humans ^1^. Azoles inhibit the sterol C14-demethylase (CYP51, also known as Erg11 in yeasts) which is one of the key enzymes in the ergosterol biosynthesis pathway ^2,3^. In most fungi, ergosterol is the most abundant sterol in the membranes, where it serves a function similar to cholesterol in human and animal cells ^4^. Inhibition of CYP51 proved to be highly effective against many fungal species, including human and animal pathogenic fungi such as dermatophytes as well as the major causative agents of invasive fungal infections such as yeasts in the genera *Cryptococcus* and *Candida* and molds in the genus *Aspergillus* ^5^. However, despite their clinical efficacy, the effects of azoles on the fungal viability and growth vary greatly among fungal species. Against certain species, especially those in the medically important genus *Candida*, they exert only a growth-inhibitory (fungistatic) activity without ultimately killing the fungal pathogen ^6^. Other pathogens, such as the medically important molds in the genus *Aspergillus* are readily killed by azoles which is referred to as fungicidal activity ^6,7^. We have previously shown that azoles trigger the formation of cell wall patches in *Aspergillus fumigatus*, the most important human pathogen of the *Aspergillus* genus. These patches are essentially cell wall bulges based on excessive synthesis of cell wall carbohydrates which extend into the lumen of the hyphae and which cause cell wall stress and contribute to the fungicidal activity of azoles against this mold ^7^. The mechanisms leading to the formation of these cell wall patches, which apparently are not formed in yeasts after azole exposure, remained unknown.

The ergosterol biosynthesis pathways of fungi differ in some parts between species. While the upstream precursors of the biosynthesis (e.g., farnesyl pyrophosphate, squalene, squalene epoxide or lanosterol) as well as the final product (ergosterol) generally appear to be the same and the biosynthesis is carried out by a highly conserved set of enzymes, the sequence of the enzymatic steps required for synthesis varies depending on the species ^8–11^. To form the initial sterols of the pathway, farnesyl pyrophosphate, the last product of the mevalonate pathway, is converted to squalene by squalene synthase (Erg9 in baker’s yeast). Squalene is then converted to squalene epoxide by squalene epoxidase (Erg1 in baker’s yeast). Squalene epoxide is converted by lanosterol synthase (Erg7 in baker’s yeast) to lanosterol, which is the first sterol of the ergosterol biosynthesis pathway ^12^.

In the baker’s yeast *Saccharomyces cerevisiae* as well as in the pathogenic yeast *Candida albicans*, lanosterol (4,4,14-trimethylcholesta-8,24(25)-dien-3β-ol (**1**); Fig. 1 A) is the substrate of CYP51 (also known as Erg11 in yeasts) which converts it to 4,4-dimethylcholesta-8,14,24(25)-trien-3β-ol (**6**). This product is then further processed by a number of other enzymes to ergosterol (**14**) ^11^. However, in the mold *A. fumigatus*, lanosterol (**1**) is converted by sterol C24-methyltransferase to eburicol (4,4,14-trimethylergosta-8,24(28)-dien-3β-ol (**2**)) ^8^. Eburicol (**2**) is then the substrate of CYP51, followed by further processing to ergosterol (**12**) ^8^. Consequently, inhibition of CYP51 results in the accumulation of different precursors in the yeasts *S. cerevisiae* and *C. albicans* on the one hand and in the mold *A. fumigatus* on the other hand.

**Fig. 1.**
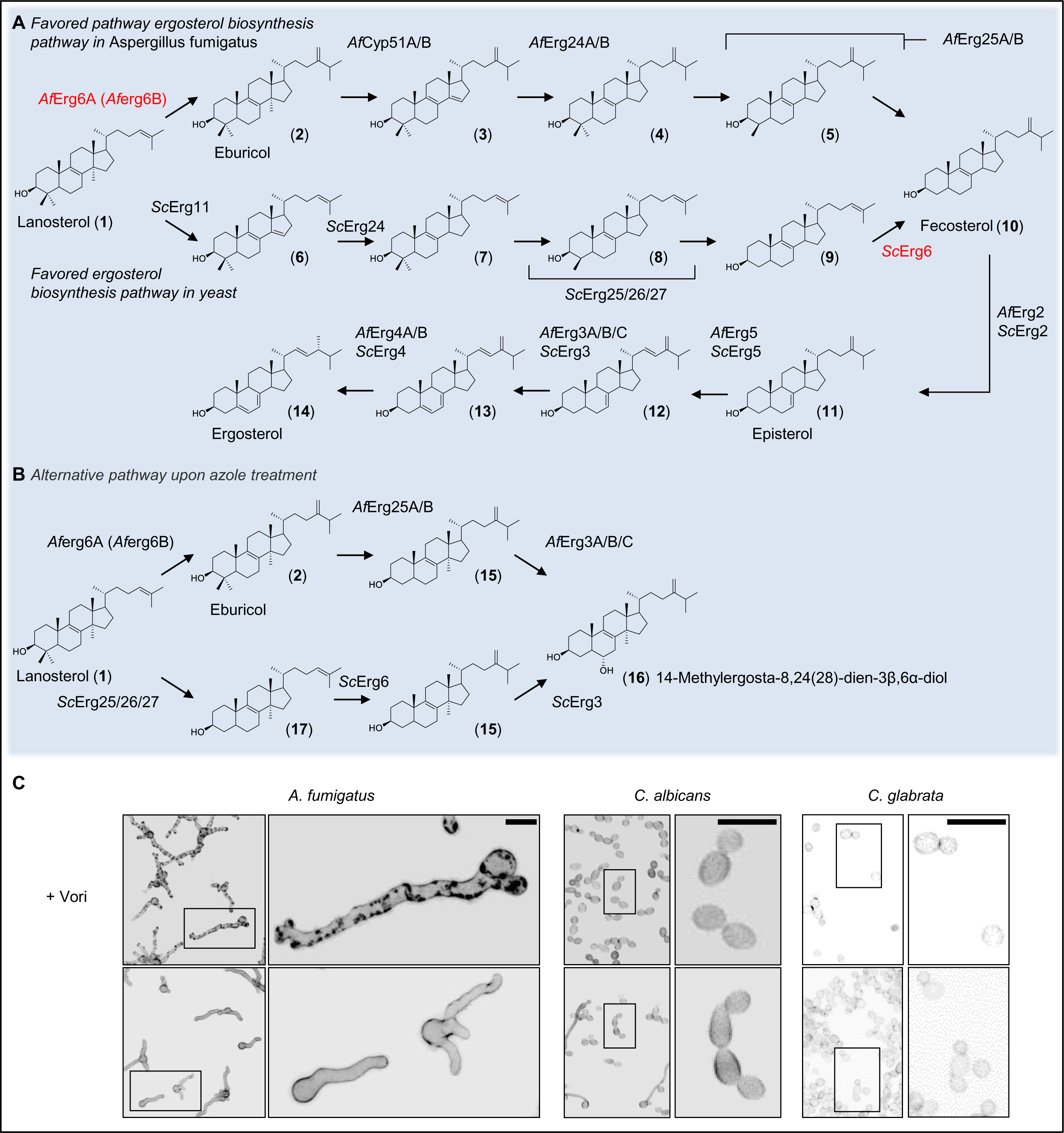
Differences in the sterol biosynthesis pathways correlate with the capability of azoles to induce cell wall carbohydrate patches in yeasts or *A. fumigatus*. (A) Ergosterol biosynthesis pathways in *A. fumigatus* (*Af,* top pathway) and *S. cerevisiae* (*Sc,* bottom pathway). In *A. fumigatus*, C24-methylation of lanosterol (**1**) by sterol C24-methyltransferase (*Af*Erg6A) is favored over C14-methylation. In yeasts, lanosterol (**1**) is the favored substrate of sterol C14-demethylase (*Sc*Erg11). Both pathways converge with the formation of fecosterol (**10**), which is then further processed to ergosterol (**14**). (B) Proposed alternative sterol biosynthesis pathway upon inhibition of sterol C14-demethylase (AfCyp51A/B and *Sc*Erg11). In yeast, lanosterol (**1**) is converted to 14-methylcholesta-8,24-dien-3β-ol (**17**), which in turn is converted to 14-methylergosta-8,24(28)-dien-3β-ol (**15**). In *A. fumigatus*, eburicol (**2**) is directly converted to 14-methylergosta-8,24(28)-dien-3β-ol (**15**). 14-methylergosta-8,24(28)-dien-3β-ol (**15**) is then converted to the 14-methylergosta-8,24(28)-dien-3β,6α-diol (**16**) which is considered a “toxic diol”. (A and B) Ergosterol biosynthesis enzymes in *A. fumigatus* (*Af*) and *S. cerevisiae* (*Sc*): sterol C24-methyltransferase (*Af*Erg6A, *Af*Erg6B and *Sc*Erg6), sterol C14-demethylase (*Af*Cyp51A/B, *Sc*Erg11), sterol C14-reductase (*Af*Erg24A/B, *Sc*Erg24), sterol C4-demethylase complex (*Af*Erg25A/B, *Sc*Erg25/26/27), sterol C8-isomerase (*Af*Erg2, *Sc*Erg2), sterol C22-desturase (*Af*Erg5, *Sc*Erg5), sterol C5-desaturase (*Af*Erg3A/B/C, *Sc*Erg3), sterol C24 reductase (*Af*Erg4A/B, *Sc*Erg4). Sterols: (**1**) lanosterol, (**2**) eburicol, (**3**) 4,4-dimethylergosta-8,14,24(28)-trien-3β-ol, (**4**) 4,4-dimethylergosta-8,24(28)-dien-3β-ol, (**5**) 4-methylergosta-8,24(28)-dien-3β-ol, (**6**) 4,4-dimethylcholesta-8,14,24-trien-3β-ol, (**7**) 4,4-dimethylcholesta-8,24-dien-3β-ol, (**8**) 4-methylcholesta-8,24-dien-3β-ol, (**9**) zymosterol, (**10**) fecosterol, (**11**) episterol, (**12**) ergosta-7,22,24(28)-trien-3β-ol, (**13**) ergosta-5,7,22,24(28)-tetraen-3β-ol, (**14**) ergosterol, (**15**) 14-methylergosta-8,24(28)-dien-3β-ol, (**16**) 14-methylergosta-8,24(28)-dien-3β,6α-diol, and (**17**) 14-methylcholesta-8,24-dien-3β-ol. (C) Conidia of *A. fumigatus* wild type and two *Candida* species (*C. albicans* ATCC14053 and C. *glabrata* ATCC2950) were inoculated in Sabouraud medium and incubated at 37 °C. After 9.5 h of incubation, the samples were either fixed and stored at 4 °C (control) or, after the medium was supplemented with 3 µg ml^-1^ voriconazole (+Vori), further incubated at 37 °C. After 15 h additional incubation, the voriconazole-exposed hyphae and yeasts were also fixed. Samples were then stained with calcofluor white and analyzed with a confocal laser scanning microscope. Depicted are representative images of z-stack projections of optical stacks of the calcofluor white fluorescence covering the entire hyphae in focus. Upper panel, voriconazole-treated hyphae; lower panel, controls. Bars represent 10 μm.

Notably, it is not in the first instance the lack of ergosterol that is responsible for the antifungal effects of azoles on yeasts. Instead, the accumulation of toxic sterols comes into play ^13^. This is best illustrated by the fact that inactivation of sterol C5-desaturase (Erg3 in baker’s yeast) causes azole resistance in yeasts. This has been shown for *C. albicans*, *Candida glabrata* and *S. cerevisiae* ^14–17^: Upon inhibition of CYP51, lanosterol (**1**) accumulates, which is converted to 14-methylfecosterol (14-methylergosta-8,24(28)-dien-3β-ol (**15**); Fig. 1 B). Erg3 then converts it to 14-methylergosta-8,24(28)-dien-3*β*,6α-diol (**16**) which is considered the “toxic diol” ^18,19^. Azole-treated yeast strains that lack the enzymatic function of Erg3 do not form the “toxic diol” (**16**). Instead, 14-methylfecosterol (**15**) accumulates, allowing them to grow (Fig. 1 B).

We hypothesized that the differences in the ergosterol biosynthesis pathways of yeasts and molds are causally involved in the different effects of azole antifungals against these fungi. To investigate the molecular nature of the different modes of action, we constructed *A. fumigatus* ergosterol biosynthesis mutants which allowed us to investigate the importance of individual ergosterol biosynthesis enzymes and dissect and assign the antimicrobial potential of individual steroid intermediates and derivatives. Our results show that it is primarily the accumulation of the ergosterol biosynthesis intermediate eburicol whose accumulation has detrimental effects on *A. fumigatus*’s cell physiology and which is responsible for the strong antimicrobial activity of azole antifungals against this pathogen.

## RESULTS

### Erg6A but not Erg6B is essential in *A. fumigatus*

Azoles induced the formation of cell wall carbohydrate patches in *A. fumigatus*, but not in pathogenic yeasts such as *C. albicans* and *C. glabrata* (Fig. 1 C). We speculated that these differences might be linked with the species-specific differences in the sequence of the enzymatic steps involved in ergosterol biosynthesis (Fig. 1 A and B).

In the first step, we aimed to generate a mutant in which the function of the sterol C24-methyltransferase (Erg6 in Baker’s yeast) can be repressed. The genome of *A. fumigatus* harbors two genes that encode Erg6 homologs, AFUA_4G03630 and AFUA_4G09190, which we named *erg6A* and *erg6B*, respectively. Since we initially assumed that the two Erg6 homologs might be functionally redundant, we planned to generate a double mutant in which one of the genes (*erg6B*) was deleted and the expression of the other gene (*erg6A*) can be downregulated by replacing the endogenous promoter with a doxycycline-inducible Tet-On promoter ^20,21^. However, we found that repression of *erg6A* alone prevents growth of *A. fumigatus*, while deletion of *erg6B* did not result in any apparent growth phenotype (Fig. 2 A). This suggested that the sterol C24-methyltransferase step, in contrast to its role in yeast ^22^, is essential for the viability of *A. fumigatus* and that *erg6B* cannot compensate for the lack of Erg6A. When comparing the growth of the conditional *erg6A_tetOn_* mutant with that of a similarly constructed conditional CYP51 *A. fumigatus* mutant (*cyp51A_tetOn_* Δ*cyp51B*) ^7^, we found that the macroscopically visible growth impairment occurred at similarly reduced doxycycline concentrations (Fig. 2 B).

**Fig. 2.**
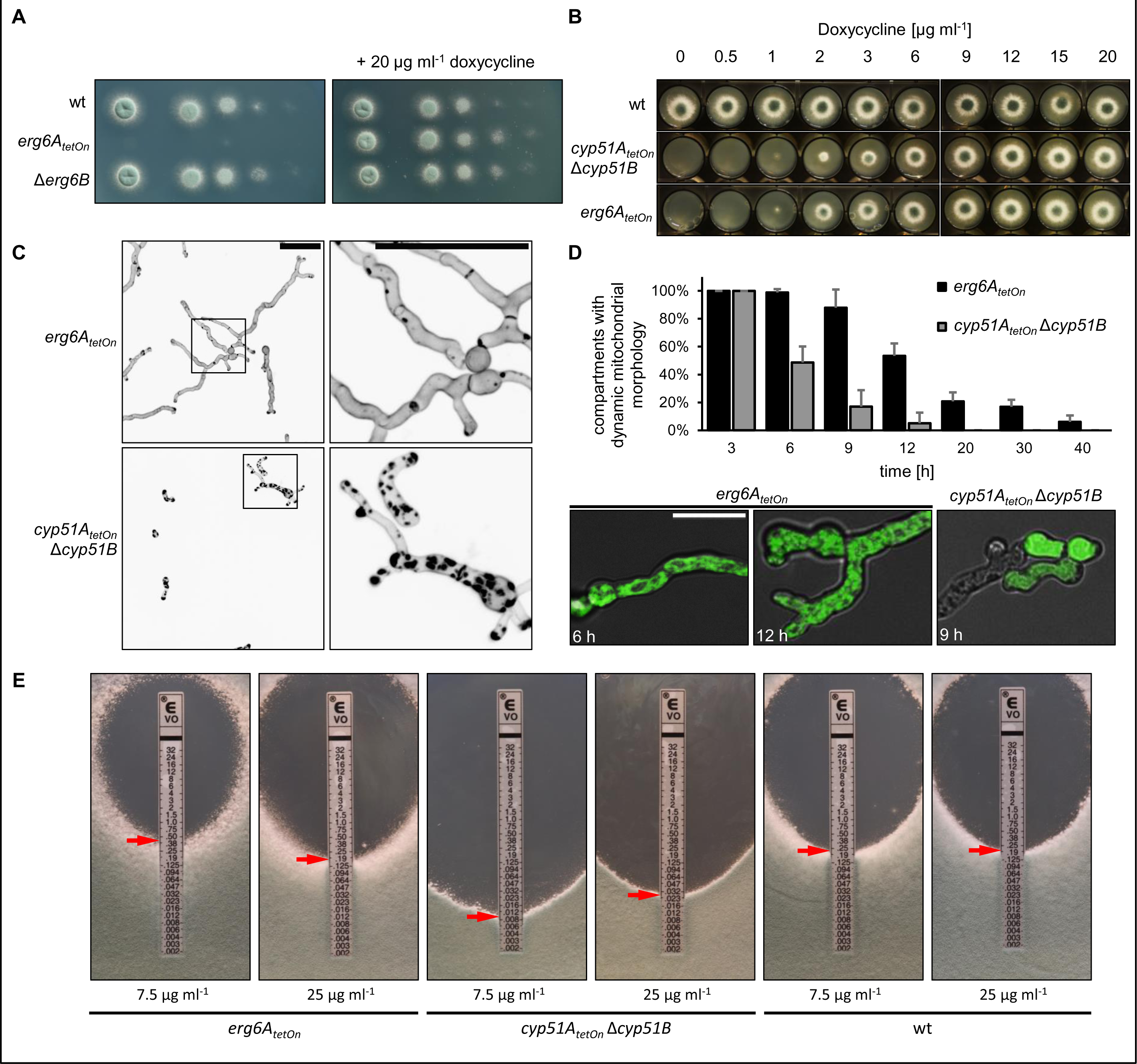
The impact of sterol C24-methyltransferase depletion on viability, cell wall patch formation and azole susceptibility. (A) In a series of 10-fold dilutions derived from a starting suspension of 5 × 10^7^ conidia ml^-1^ of wild type (wt), the conditional mutant *erg6A_tetOn_*, and the deletion mutant Δ*erg6B*, aliquots of 3 µl were spotted onto AMM agar plates. AMM was supplemented with 20 μg ml^−1^ doxycycline when indicated. Agar plates were incubated at 37 °C, representative photos were taken after 28 h. (B) 1.5 × 10^3^ conidia of the indicated strains were spotted on AMM agar supplemented with the indicated amount of doxycycline. The plates were then incubated at 37 °C, representative photos were taken after 40 h. (C and D) Conidia of the *erg6A_tetOn_* and *cyp51A_tetOn_* Δ*cyp51B* mutants which express mitochondria-targeted GFP (D) or not (C) were inoculated in Sabouraud medium supplemented with 20 µg mL^-1^ doxycycline and incubated at 37 °C. After 6 h, doxycycline was depleted by washing the wells three times with Sabouraud medium without doxycycline. The hyphae were then incubated in Sabouraud medium without doxycycline at 37 °C for an additional 40 h. (C) Hyphae were stained with calcofluor white and analyzed with a confocal microscope. Depicted are representative images of z-stack projections of optical stacks of the calcofluor white fluorescence covering the entire hyphae in focus. The right images show magnifications of the framed sections in the left images. Bars represent 50 μm and are applicable to all images in the respective panel. (D, column graph) At the indicated time points after doxycycline depletion, the viability of the hyphae of the indicated strains expressing mitochondria-targeted GFP was analyzed with time-lapse spinning disc confocal microscopy. The bars indicate the percentage of hyphal compartments with evident mitochondrial dynamics (viable compartments). Data points represent the means of five technical replicates for each timepoint, with an average of approx. 80 analyzed compartments per strain for each timepoint. The error bars indicate standard deviations based on the five technical replicates. (D, lower panel) Representative overlay images of the bright field channel and of z-stack projections of optical stacks of the GFP fluorescence covering the entire hyphae in focus. Depicted are exemplary images of a viable hypha with tubular mitochondrial morphology (also showing mitochondrial dynamics in time-lapse microscopy; left image) and of hyphae with fragmented mitochondria (showing no mitochondrial dynamics in time-lapse microscopy; middle image) or with no or cytosolic GFP signal (right image) which were considered to be dead. Bars represent 50 μm and are applicable to all respective subpanels. (E) 1 × 10^6^ conidia of the indicated strains were spread on Sabouraud agar plates. Agar was supplemented with the indicated amount of doxycycline to achieve a different induction of the conditional promoters. Voriconazole Etest strips were applied. The plates were incubated at 37 °C and representative photos were taken after 42 h.

### Depletion of CYP51 but not of Erg6A triggers cell wall patch formation

We next analyzed whether repression of *erg6A* results in the formation of cell wall carbohydrate patches, similar as we had previously found it in *A. fumigatus* that was treated with azoles or where CYP51 was depleted ^7^. As shown in Fig. 2 C, depletion of CYP51 resulted in the accumulation of chitin-rich calcofluor white-stainable cell wall patches. In contrast, depletion of Erg6A did not result in the accumulation of cell wall patches (Fig. 2 C). Notably, compared to hyphae of the conditional CYP51 mutant, the hyphae of the conditional *erg6A_tetOn_* mutant continued to grow for much longer after the removal of doxycycline. We speculated that the time of survival of the *erg6A_tetOn_* mutant after doxycycline depletion also differs from that of the CYP51 mutant after doxycycline depletion. This could potentially correlate with a delay in formation of cell wall patches. To study this, we used an assay which is based on the evaluation of mitochondrial dynamics to assess the viability of *Aspergillus* hyphae over time ^7^. As shown in Fig. 2 D, the half life of *erg6A_tetOn_* mutant hyphae after doxycycline depletion was approximately double that of the CYP51 mutant hyphae after doxycycline depletion. Since the photos depicted in Fig. 2 C were taken after a similar incubation time (40 hours) under similar experimental conditions, this indicates that the *erg6A_tetOn_* mutant died without forming cell wall patches prior to death.

### Depletion of Erg6A increases azole tolerance of *A. fumigatus*

Our results demonstrated that Erg6A is essential for viability of *A. fumigatus*. But our results also suggested that the cell biological nature of Erg6A depletion-induced death is somewhat different from that of the CYP51 depletion- or inhibition-induced death because 1) death occurs much slower and 2) no cell wall patches get formed prior to death. We speculated that the accumulation of different ergosterol precursors could be linked with this. Erg6A depletion would presumably result in accumulation of lanosterol (**1**), CYP51 depletion results in accumulation of eburicol (**2**) according to the literature (Fig. 1 A) ^23^. This would be compatible with a model where the accumulation of eburicol is more toxic than the accumulation of lanosterol.

We then tested whether Erg6A depletion impacts on the azole susceptibility of *A. fumigatus*, and compared this to the effects of depletion of CYP51 on azole sensitivity. Reduced expression of CYP51 resulted in increased susceptibility to azoles (Fig. 2 E), which is perfectly in line with overexpression of CYP51 having the opposite effect and being a major resistance mechanism against azoles in *A. fumigatus* ^24^. Interestingly, culturing the conditional *erg6A_tetOn_* mutant under less inducing conditions resulted in reduced susceptibility to azoles when compared to wild type and the *erg6A_tetOn_* mutant under more inducing conditions (Fig. 2 E). This indicates that a reduced synthesis of eburicol is beneficial for *A. fumigatus* if CYP51 is inhibited by azoles at the same time.

### Erg1 and Erg9 are essential, but their depletion or inhibition does not result in excessive cell wall patch formation

To confirm that the azole-induced cell wall carbohydrate patches are not just a result of reduced sterol biosynthesis, we constructed conditional squalene synthase (Erg9) and squalene epoxidase (Erg1) mutants ^25^. Erg9 and Erg1 are both essentially required to form squalene or, respectively, squalene epoxide which are the precursors of lanosterol, the first sterol in the ergosterol biosynthesis pathway ^12^. Consequently, downregulation of *erg9* or *erg1* should result in disruption of the whole sterol biosynthesis. The conditional *erg9_tetOn_* and *erg1_tetOn_* mutants were not viable under repressed conditions (Fig. 3 A) ^25^. Interestingly, none of these mutants formed cell wall patches under repressed conditions (Fig. 3 B). Similarly, *A. fumigatus* wild type, which was expressing mitochondria-targeted green fluorescent protein (GFP) as viability marker, did not form cell wall patches when treated with the Erg1 inhibitor terbinafine (Fig. 3 C). The mitochondrial dynamics-based viability marker demonstrated that the terbinafine-treated hyphae had died without forming patches (Fig. 3 C). This demonstrates that disruption of the whole sterol biosynthesis is lethal, but does not result in the formation of cell wall patches.

**Fig. 3.**
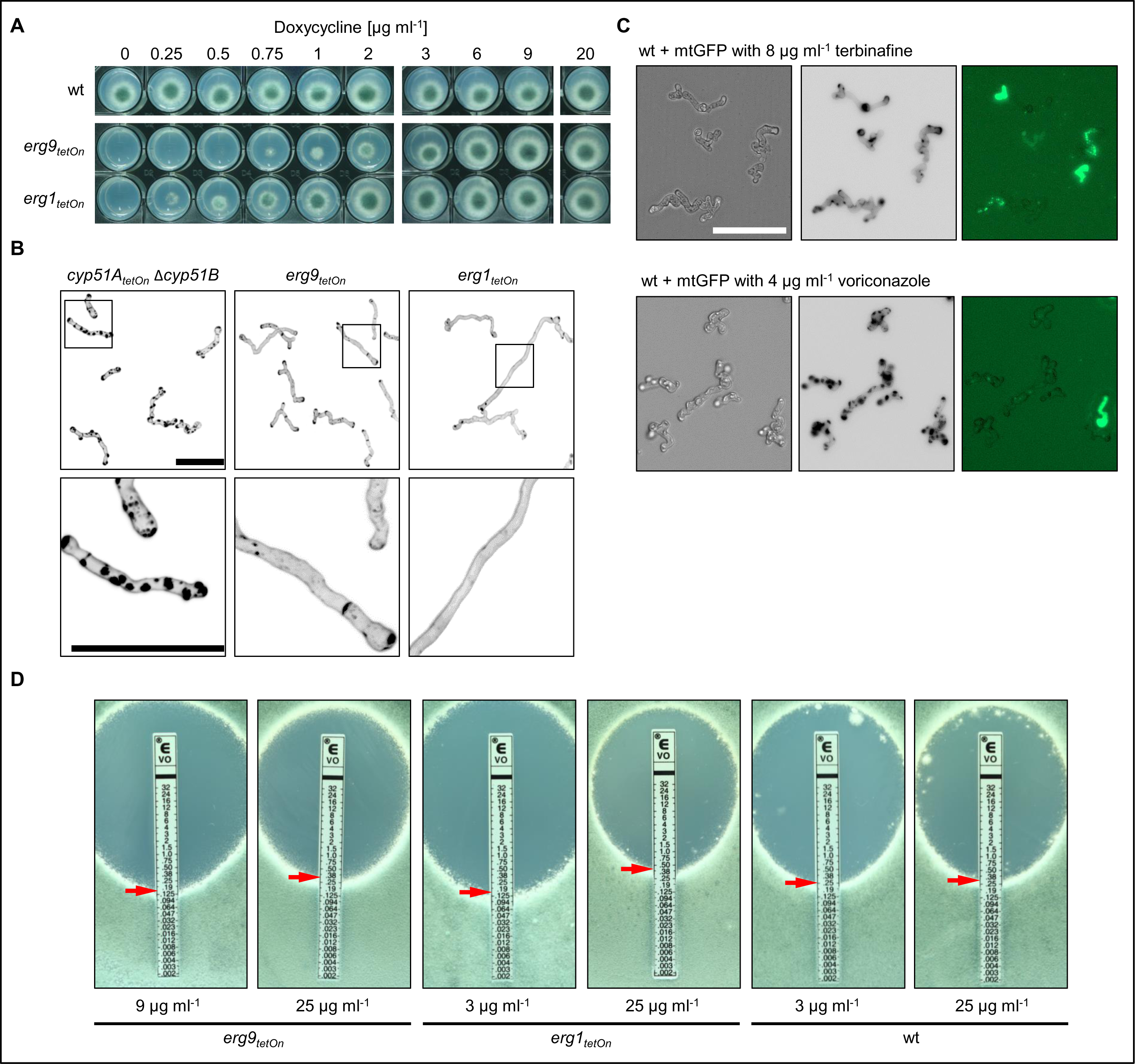
Depletion of the squalene epoxidase or the squalene synthase does not trigger excessive cell wall patch formation, but their altered expression results in changes in azole sensitivity. (A) 1.5 × 10^3^ conidia of wild type (wt) or of the conditional squalene epoxidase (*erg9_tetOn_*) or of the squalene synthase (*erg1_tetOn_*) mutants were spotted on AMM agar supplemented with the indicated amount of doxycycline. The plates were then incubated at 37 °C, representative photos were taken after 40 h. (B) Conidia of the indicated strains were inoculated in Sabouraud medium supplemented with 20 µg ml^-1^ doxycycline and incubated at 37 °C. After 6 h, doxycycline was depleted by washing the wells three times with Sabouraud medium without doxycycline. The hyphae were then incubated in Sabouraud medium without doxycycline at 37 °C for an additional 40 h and subsequently stained with calcofluor white and analyzed with a confocal microscope. Depicted are representative images of z-stack projections of optical stacks of the calcofluor white fluorescence covering the entire hyphae in focus. The lower images show magnifications of the framed sections in the upper images. Bars represent 50 μm and are applicable to all images in the respective panel. (C) Conidia of wild-type expressing mitochondria-targeted GFP were inoculated in Sabouraud medium. After 8 h incubation at 37 °C, medium supplemented with 8 µg ml^-1^ terbinafine (upper panel) or with 4 µg ml^-1^ voriconazole (lower panel). After an additional 16 h incubation at 37 °C, hyphae were analyzed with a fluorescence microscope. Fluorescence signals were analyzed sequentially, first the GFP signal was recorded followed by staining with calcofluor white and recording of the calcofluor white signal. Depicted are representative images of bright-field (left) and z-stack projections of optical stacks of the calcofluor white fluorescence (middle) and GFP fluorescence (right) after deconvolution, that cover the entire hyphae in focus. The bar represents 50 µm and is applicable to all images. (E) 1 × 10^6^ conidia of the indicated strains were spread on Sabouraud agar plates. Agar was supplemented with the indicated amount of doxycycline to achieve a different induction of the conditional promoters. Voriconazole Etest strips were applied. The plates were incubated at 37 °C and representative photos were taken after 42 h.

Next, we analyzed the impact of reduced *erg9* and *erg1* expression on azole susceptibility. As shown in Fig. 3 D, less inducing conditions resulted in increased azole susceptibility of the conditional *erg9_tetOn_* and *erg1_tetOn_* mutants compared to wild type. In contrast, more inducing conditions resulted in an azole susceptibility which was similar compared to wild type (Fig. 3 D). Taken together, these findings show that azoles are more effective when the total sterol biosynthesis is downregulated. Conversely, a presumable upregulation of total sterol biosynthesis results in a lower susceptibility to azoles.

### Impact of Erg9, Erg6A and CYP51 depletion on fungal sterol patterns

The conclusions presented above were based on the assumption that in *A. fumigatus* depletion of CYP51, similar to treatment with azole antifungals, would result in accumulation of eburicol (**2**), and depletion of Erg6A would result in accumulation of lanosterol (**1**). Furthermore, we would not expect a drastic change in the sterol pattern upon depletion of Erg9 or Erg1 because any synthesized lanosterol could still be readily converted to ergosterol (**14**). To confirm these suppositions, we analyzed the sterol patterns of the conditional CYP51, *erg6_tetOn_* and *erg9_tetOn_* mutants under inducing as well as repressing conditions (Fig. 4 A). Different concentrations of doxycycline, the chemical used to induce the conditional TetOn promoters, did not significantly alter the sterol pattern of wild type (Supplementary Fig. 1 A). When compared with wild type, depletion of Erg9 resulted in a slight but significant decrease in the amount of ergosterol relative to other sterols (Fig. 4 B, Supplementary Fig. 1 B). Further depletion of Erg9 (by further reducing the doxycycline concentration) did not result in a remarkable alteration of this pattern (Fig. 4 B, Supplementary Fig. 1 B). Depletion of Erg6 resulted in a significant decrease of ergosterol and a relative increase of lanosterol (Fig. 4 B, Supplementary Fig. 1 B). Depletion of CYP51 resulted in a dramatic increase of eburicol (**2**), but also in a remarkable increase of the presumably “toxic diol” (14-methylergosta-8,24(28)-dien-3β,6α-diol (**16**)), and lanosterol (**1**). The impact of low doxycycline concentrations on the sterol patterns was much more pronounced in the conditional CYP51 mutant compared to the conditional *erg6_tetOn_* mutant (Supplementary Fig. 1 B). We speculated that this could be linked with the complete lack of *cyp51B* in the CYP51 mutant (*cyp51A_tetOn_* Δ*cyp51B*) and the inability of the Tet-On promoter-dependent *cyp51A* to compensate for this (see also increased azole susceptibility under fully induced conditions; Fig. 2 E).

**Fig. 4.**
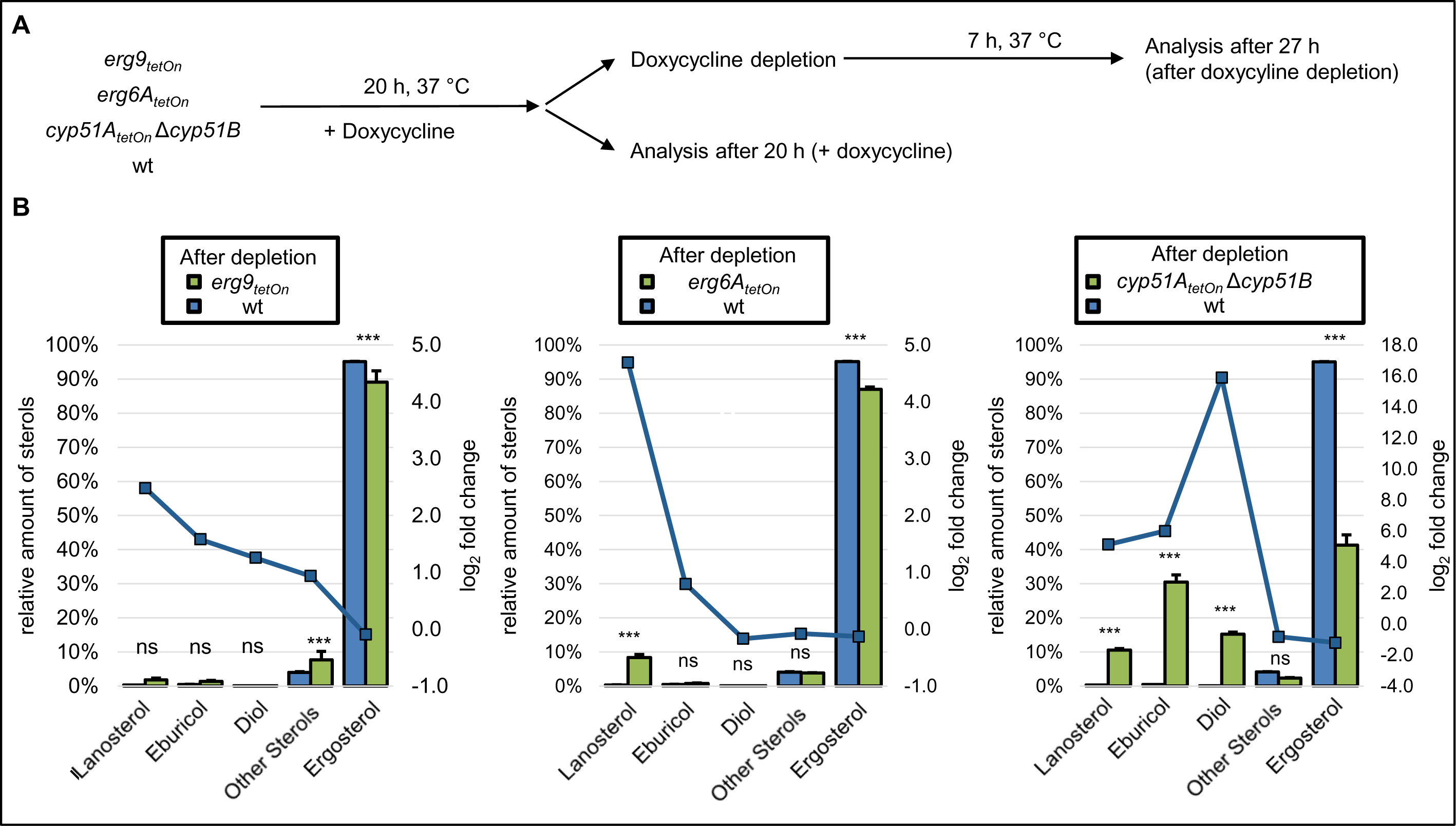
Sterol profiles of squalene synthase-, sterol C24-methyltransferase- and sterol C14-demethylase-depleted *A. fumigatus* hyphae. (A) Conidia of wild type (wt) and the indicated mutants were inoculated in Sabouraud liquid medium. Cultures with the conditional sterol C24-methyltransferase mutant (*erg6A_tetOn_*), the conditional sterol C14-demethylase mutant (*cyp51A_tetOn_* Δ*cyp51B*) and the respective wild-type control (wt) were supplemented with 2 µg ml^-1^ doxycycline. Cultures with the conditional squalene synthase (*erg9_tetOn_*) and the respective wild-type control (wt) were supplemented with 3 µg ml^-1^ doxycycline. After 20 h incubation in a rotary shake at 37 °C, mycelium was either directly harvested or washed with and transferred into Sabouraud liquid medium without doxycycline, incubated in a rotary shake at 37 °C for another 7h, and then harvested. For each condition, three biological replicates were cultured. (B) The sterol patterns of the harvested mycelia were analyzed by gas chromatography-mass spectrometry (GC-MS). The column graphs show the relative amounts (percentage of total sterol, left y-axis) of the indicated sterols for the indicated strains before and after doxycycline depletion. The data points with the square symbols indicate the log_2_-fold change (right y-axis) in the amount of the respective sterol of the pairwise comparison of the conditions shown in the individual graphs. The log_2_-fold change data points were connected by lines for better visual illustration of the changes in the profiles. The column bar data points represent means of three replicates per condition, the error bars indicate standard deviations. Statistical significance (* p<0.05; ** p<0.01; *** p<0.001) was calculated with a two-way ANOVA with Tukey’s multiple comparison test. Further pairwise comparisons of the results of this experiment are shown in Supplementary Fig. 1.

These findings confirmed our expectation that Erg6 depletion results in accumulation of lanosterol. They also confirmed the previously reported accumulation of eburicol upon depletion of CYP51 ^23,26^. However, our findings also demonstrate that the presumably “toxic diol” (**15**) is formed in *A. fumigatus* when CYP51 functionality is lacking.

### ERG3-depleted hyphae form cell wall patches when exposed to azoles

Depletion of CYP51 resulted in a significant increase of the presumably “toxic diol” (**16**). Accumulation of this sterol is responsible for the antifungal effect of azole antifungals against yeasts ^13^. Accordingly, its accumulation might also explain the antifungal effects of azoles on *A. fumigatus*, e.g., the formation of cell wall carbohydrate patches. The formation of the “toxic diol” in yeasts requires the incomplete (defective) reaction of the sterol C5-desaturase. If this model also holds true in *Aspergillus*, disruption of the desaturation step of 14-methylfecosterol (**15**) by inactivating sterol C5-desaturase should cause azole resistance in A. fumigatus, similar as it was previously shown for S. cerevisiae and pathogenic yeasts 14,15,17. The genome of A. fumigatus harbors three genes that encode homologes of S. cerevisiae Erg3, AFUA_6G05140 (erg3A), AFUA_2B00320 (erg3B), and AFUA_8G01070 (*erg3C*). While mutants lacking the individual enzymes as well as an *erg3A erg3B* double mutant have been reported and characterized previously ^8,27^, no mutants that lack the other combinations (*erg3A erg3C* and *erg3B erg3C*) or all three enzymes have been reported. To characterize the roles of Erg3A, Erg3B, and Erg3C, we constructed single, double and triple mutants (Fig. 5 A and B). As shown in Fig. 5 B, the lack of individual Erg3 enzymes as well as the lack of Erg3A and Erg3B or Erg3B and Erg3C did not result in any significant impact on growth. In contrast, the lack of Erg3A and Erg3C as well as the lack of Erg3A, Erg3B and Erg3C resulted in reduced growth rates and a reduced production of asexual spores (conidia) (Fig. 5 B).

**Fig. 5.**
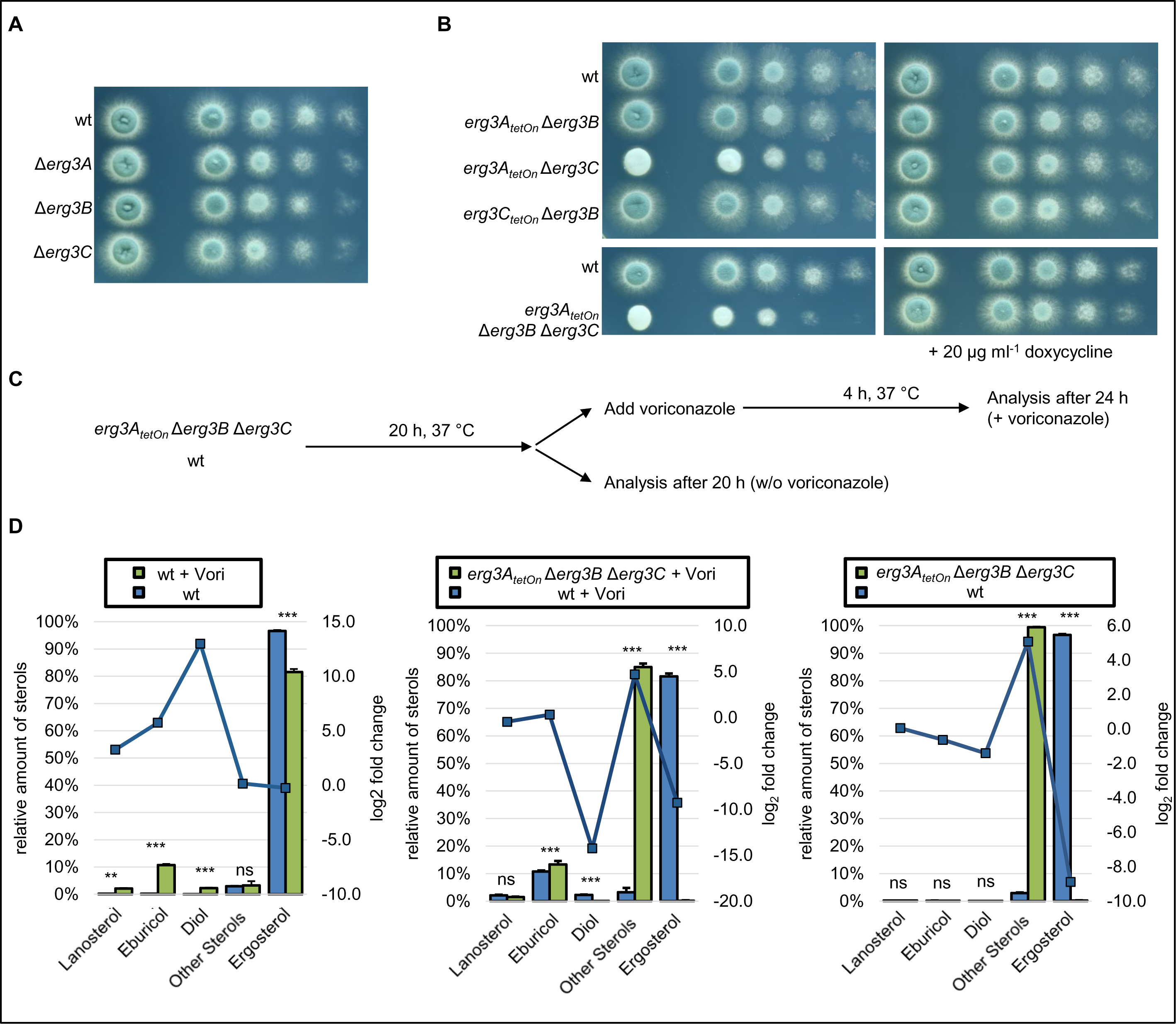
Construction of a conditional sterol C5-desaturase mutant repression and the impact of sterol C5-desaturase depletion on the sterol pattern of azole-treated hyphae. (A and B) In a series of 10-fold dilutions derived from a starting suspension of 5 × 10^7^ conidia ml^-1^ of wild type (wt) and the indicated sterol C5-desaturase mutant single, double and triple mutants, aliquots of 3 µl were spotted onto AMM agar plates. AMM was supplemented with 20 μg ml^−1^ doxycycline when indicated. Agar plates were incubated at 37 °C, representative photos were taken after 28 h. (C and D) Conidia of wild type and the conditional sterol C5-desaturase mutant (*erg3A_tetOn_* Δ*erg3B* Δ*erg3C*) were inoculated in Sabouraud medium. After 20 h incubation in a rotary shake at 37 °C, mycelium was either directly harvested or supplemented with 2 µg ml^-1^ voriconazole (+Vori), incubated in a rotary shake at 37 °C for another 4 h, and then harvested. For each condition, three biological replicates were cultured. (D) The sterol patterns of the harvested mycelia were analyzed by gas chromatography-mass spectrometry (GC-MS). The column graphs show the relative amounts (percentage of total sterol, left y-axis) of the indicated sterols for the indicated strains under the indicated conditions. The data points with the square symbols indicate the log_2_-fold change (right y-axis) in the amount of the respective sterol of the pairwise comparison of the conditions shown in the individual graphs. The log_2_-fold change data points were connected by lines for better visual illustration of the changes in the profiles. The column bar data points represent means of three replicates per condition, the error bars indicate standard deviations. Statistical significance (* p<0.05; ** p<0.01; *** p<0.001) was calculated with a two-way ANOVA with Tukey’s multiple comparison test.

We then asked whether the conditional ERG3 triple mutant (*erg3A_tetOn_* Δ*erg3B* Δ*erg3C*) still forms the presumably “toxic diol” upon exposure to azoles. To this end, we analyzed the sterol profiles of the repressed conditional ERG3 mutant in the absence and presence of azoles and compared them with the sterol profiles of wild type cultured under similar conditions (Fig. 5 C and D). As expected, the ERG3 mutant under repressed conditions did not form ergosterol (**14**) because of the lack of sterol C5-desaturase function, but instead accumulated Δ^7^-sterols summed as “other sterols” (relative percentages of total sterols: 74 % ergosta-7,22,24(28)-trien-3β-ol (**12**), 15 % ergosta,7,24(28)-dien-3β-ol (episterol, **11**), 9 % ergosta-7,22-dien-3β-ol (5-dihydroergosterol); Fig. 5 D). The azole-treated wild type accumulated eburicol, lanosterol and the presumably “toxic diol” with a concomitant relative decrease of ergosterol (Fig. 5 D). While the azole-exposed ERG3 mutant under repressed conditions also accumulated eburicol, lanosterol and other sterols (relative percentages of total sterols: 66 % ergosta-7,22,24(28)-trien-3β-ol (**12**), 10 % ergosta-7,22-dien-3β-ol (5-dihydroergosterol), 8 % ergosta,7,24(28)-dien-3β-ol (episterol, **11**)), it did not form the presumably toxic diol (Fig. 5 D). This confirms that the Erg3 enzymes in *A. fumigatus* are needed to form 14-methylergosta-8,24(28)-dien-3β,6α-diol (“toxic diol” (**16**)) upon exposure to azoles.

We next evaluated whether the formation of the “toxic diol” (**16**) is required for the formation of the azole-induced cell wall patches. To this end, we exposed hyphae of the ERG3 mutant under repressed conditions to azoles. As shown in Fig. 6 A, the lack of ERG3 did not suppress the formation of cell wall patches after exposure to azole antifungals. This demonstrates that it is not the accumulation of the “toxic diol” which triggers the formation of the azole-induced cell wall patches in *A. fumigatus*.

**Fig. 6.**
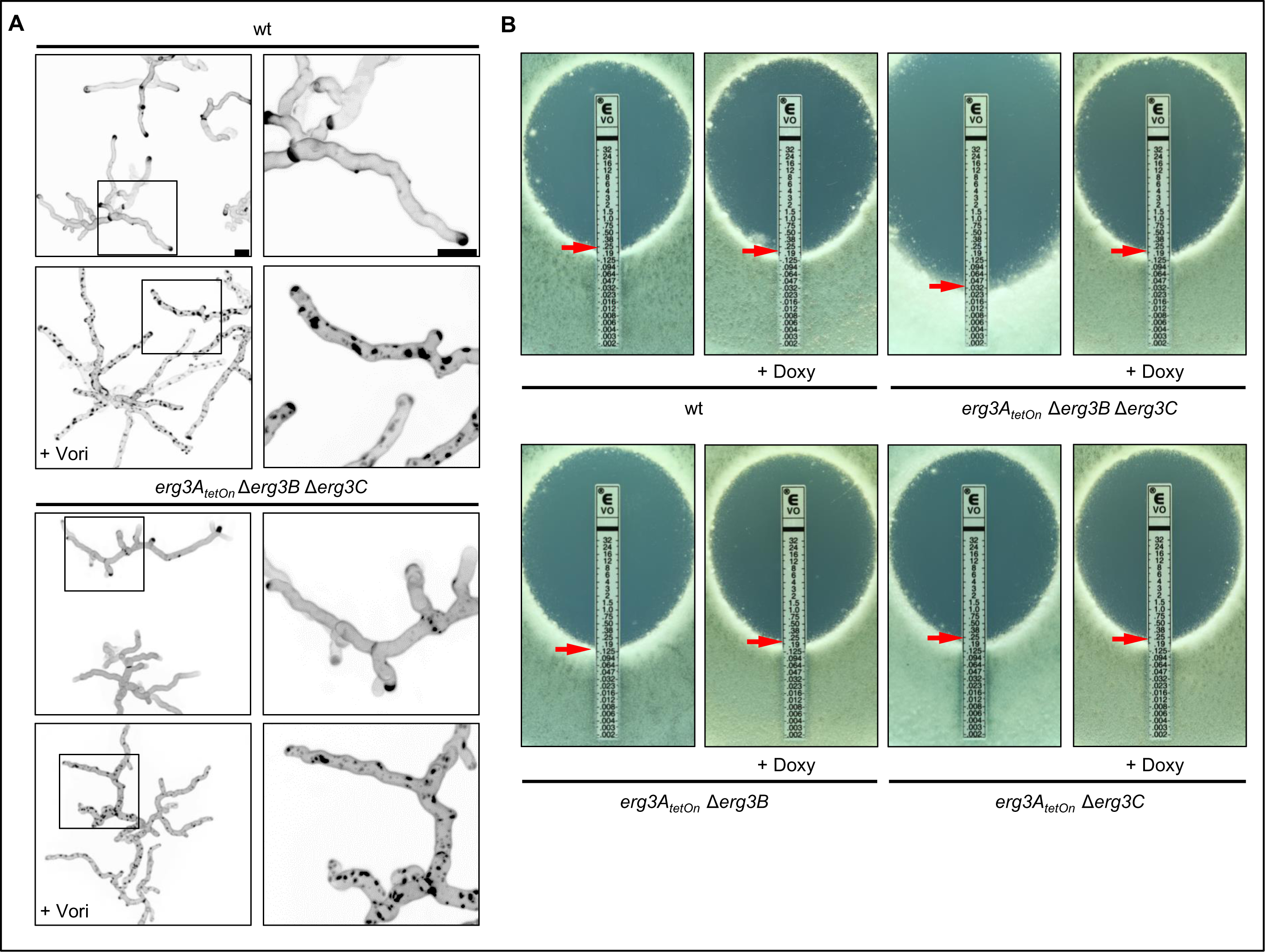
The influence of the sterol C5-desaturase on azole susceptibility and azole-induced cell wall patch formation. (B) 1 × 10^6^ conidia of wild type (wt) and the indicated strains were spread on Sabouraud agar plates. When indicated, agar was supplemented with 25 μg ml^−1^ doxycycline. Voriconazole Etest strips were applied. The plates were incubated at 37 °C and representative photos were taken after 42 h. (A) Conidia of wild type and the conditional sterol C5-desaturase mutant (*erg3A_tetOn_* Δ*erg3B* Δ*erg3C*) were inoculated in Sabouraud medium. After 10 h, the samples were either directly analyzed (controls without azole) or the medium was supplemented with 4 µg ml^-1^ voriconazole (+Vori). The voriconazole-exposed hyphae were then incubated at 37 °C for another 5 h. The untreated hyphae (upper images per indicated strain) and the voriconazole-treated hyphae (lower images per indicated strain) were stained with calcofluor white and analyzed with a confocal microscope. Depicted are representative images of z-stack projections of optical stacks of the calcofluor white fluorescence covering the entire hyphae in focus. The right panels show magnifications of the framed sections in the left panels. Bars represent 50 μm and are applicable to all images in the respective panel.

### Depletion of the ERG3 decreases azole tolerance of *A. fumigatus*

In yeast, the lack of Erg3 results in azole resistance ^13,18,19^. We asked whether this also holds true in *A. fumigatus*. Growth tests were performed to analyze the azole susceptibilities of the conditional Erg3 double mutants and the conditional ERG3 triple mutant under induced and repressed conditions. As shown in Fig. 6 B, the azole susceptibilities of the *erg3A_tetOn_* Δ*erg3B* mutant and the *erg3A_tetOn_* Δ*erg3C* mutant were not significantly changed under induced or repressed conditions when compared with wild type. Similarly, the ERG3 triple mutant under induced, effectively representing a mutant lacking Erg3B and Erg3C, showed no significantly altered azole susceptibility (Fig. 6 B). However, under repressed conditions, the ERG3 triple mutant was significantly more susceptible to azoles (Fig. 6 B). This shows that Erg3A/B/C are at least partially functionally redundant and counteract the toxic effects of azole antifungals in *A. fumigatus*. This is the exact opposite of what has been described for the mode of action of azoles and the role of Erg3 therein for yeasts.

### Disruption of mitochondrial complex III in azole–treated *A. fumigatus* enhances conversion of eburicol to 14-methylergosta-8,24(28)-dien-3β,6α-diol

We have previously shown that mutants that lack a functional mitochondrial complex III show an unusual azole susceptibility phenotype ^7,28^. Lack of a functional complex III results in a significantly reduced minimal inhibitory concentration (MIC) of azole antifungals. At the same time, the complex III mutants were able to survive at azole concentrations above the MIC, which is in marked contrast to the wild type which is effectively killed. This correlates with a decrease in azole-induced cell wall patches ^7^. A possible explanation for the unusual sensitivity to azoles could be related to an altered processing of sterols. To explore this, the sterol profile of a conditional cytochrome *c* mutant (*cycA_tetOn_*) and a conditional Rieske protein mutant (*rip1_tetOn_*) were analyzed and compared to wild type (Fig. 7 A). Under repressed conditions, the sterol profiles of the *cycA_tetOn_* and *rip1_tetOn_* mutants were similar to that of wild type (Fig. 7 B). After exposure to azoles, the ergosterol levels decreased more rapidly in the *cycA_tetOn_* and *rip1_tetOn_* mutant than in the wild type (Fig. 7 C). Surprisingly, significantly more eburicol accumulated in the azole-treated wild type than in the azole-treated repressed *cycA_tetOn_* and *rip1_tetOn_* mutants, whereas significantly more “toxic diol” (14-methylergosta-8,24(28)-dien-3β,6α-diol (**16**)) accumulated in the the *cycA_tetOn_* and *rip1_tetOn_* mutants under repressed conditions than in the wild type (Fig. 7 C). This demonstrates that mutants that lack mitochondrial complex III functionality convert eburicol much more efficiently into the diol than the wild type.

**Fig. 7.**
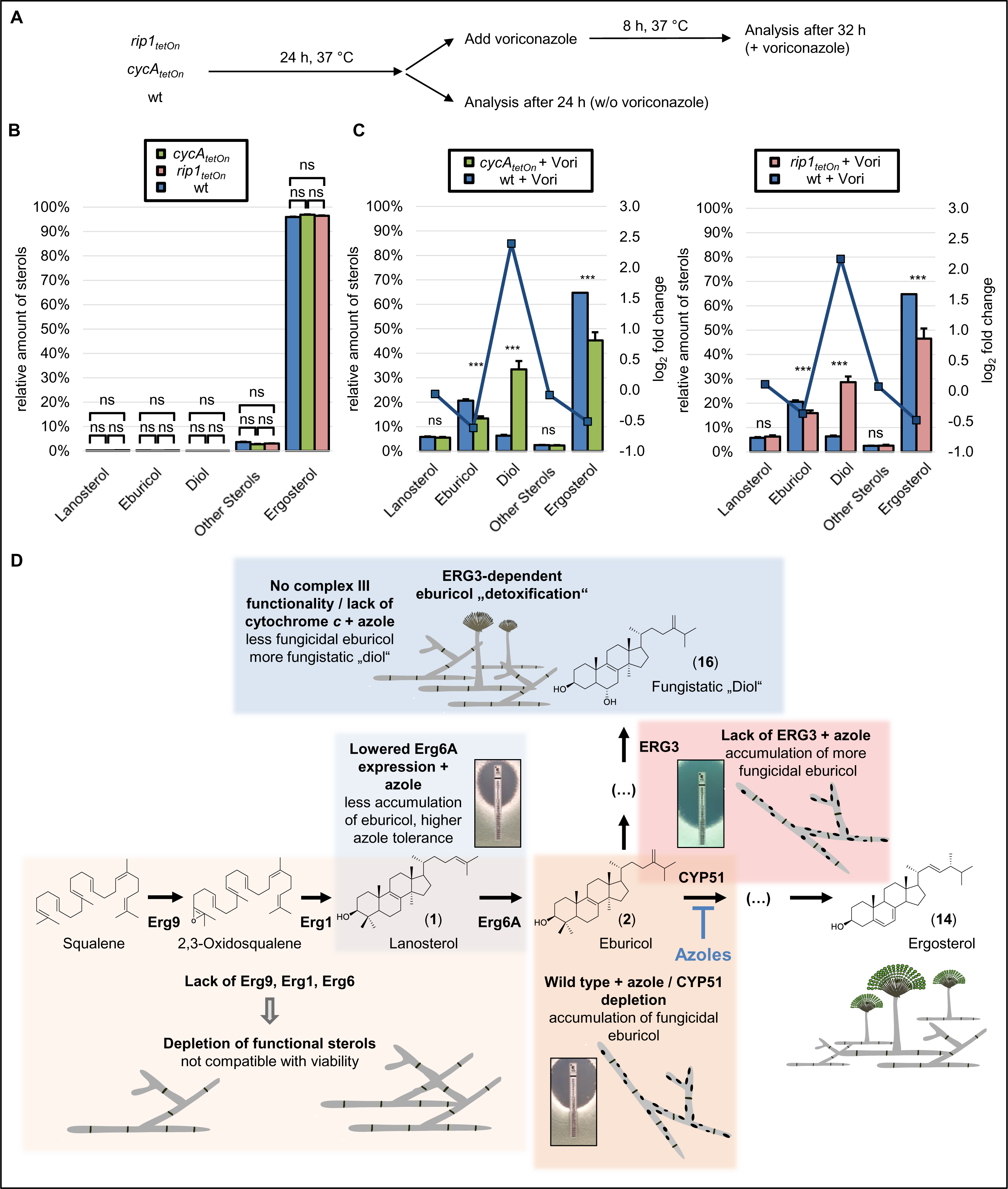
Increased formation of 14-methylergosta-8,24(28)-dien-3β,6α-diol in cytochrome *c*-depleted hyphae and model of the antifungal activity of azoles against *A. fumigatus*. (A - C) Conidia of wild type (wt) and conditional mutants lacking cytochrome *c* (*cycA_tetOn_*) or the Rieske protein (*rip1_tetOn_*) in the absence of doxycycline were inoculated in Sabouraud liquid medium. After 24 h incubation in a rotary shake at 37 °C, mycelium was either directly harvested or supplemented with 1 µg ml^-1^ voriconazole (+Vori), incubated in a rotary shake at 37 °C for another 8 h, and then harvested. For each condition, three biological replicates were cultured. (B and C) The sterol patterns of the harvested mycelia were analyzed by gas chromatography-mass spectrometry (GC-MS). The column graphs show the relative amounts (percentage of total sterol, left y-axis) of the indicated sterols for the indicated strains without voriconazole (B) or after exposure to voriconazole (C). The data points with the square symbols indicate the log_2_-fold change (right y-axis) in the amount of the respective sterol of the pairwise comparison of the conditions shown in the individual graphs. The log_2_-fold change data points were connected by lines for better visual illustration of the changes in the profiles. The column bar data points represent means of three replicates per condition, the error bars indicate standard deviations. Statistical significance (* p<0.05; ** p<0.01; *** p<0.001) was calculated with a two-way ANOVA with Tukey’s multiple comparison test. (D) Model of the mode of action of azole antifungals in the pathogenic mold *A. fumigatus*. Azole antifungals inhibit sterol C14-demethylase (CYP51). This results in accumulation of eburicol (**2**) which is fungicidal and triggers the formation of cell wall carbohydrate patches. A lack of squalene synthase (Erg9), squalene epoxidase (Erg1) or sterol C24-methyltransferase (Erg6A) results in depletion of functional sterols, thus in suppression *A. fumigatus’* growth. However, this depletion does not result in the formation of the fungicidal cell wall carbohydrate patches. Lowered expression of Erg6A results in increased azole resistance because of reduced formation and, following azole treatment, reduced accumulation of eburicol. Following azole exposure, the accumulating eburicol is partially converted into the less toxic, fungistatic “diol” (14-methylergosta-8,24(28)-dien-3*β*,6α-diol; **16**). While in *A. fumigatus* wild type this conversion is less efficient, it is increased in *A. fumigatus* mutants that lack a functional mitochondrial complex III, explaining the fungistatic activity of azoles against and the reduced formation of cell wall carbohydrate patches in these mutants. A lack of sterol C5-desaturase (Erg3) does not result in increased resistance, but causes increased susceptibility to azole antifungals since the accumulating eburicol cannot be converted into the less toxic, fungistatic “diol”.

## DISCUSSION

Azole antifungals are the mainstay for the treatment and prophylaxis of fungal infections. In yeasts such as *C. albicans* and *C. glabrata*, azole treatment results in the accumulation of lanosterol (**1**), which is converted to 14-methylergosta-8,24(28)-dien-3β,6α-diol (**16**), the so-called “toxic diol”, whose buildup is responsible for the antifungal effect ^13^. However, our present study demonstrates that treatment of *A. fumigatus* with azoles results in a predominant accumulation of eburicol (**2**), but also of lanosterol and the “toxic diol”, which is in agreement with previous studies ^23,26^. The impact of these differences remained unknown. While the azole activity against yeasts is primarily fungistatic, this drug class exerts a fungicidal effect against molds ^6,7^. We have previously shown that azole-induced cell wall carbohydrate patches contribute to the fungicidal effect of azoles against *A. fumigatus* ^7^. Based on these findings, we speculated that the eburicol accumulation could contribute to the formation of cell wall carbohydrate patches. Remarkably, in the present study we were able to show that the lack of other ergosterol biosynthesis enzymes, such as sterol C24-methyltransferase (Erg6A), squalene synthase (Erg9) and squalene epoxidase (Erg1), which are also essential for the viability of *A. fumigatus*, did not trigger the formation of cell wall carbohydrate patches. This demonstrates that the cell wall carbohydrate patch formation is specifically linked with the inhibition of sterol C14-demethylase (CYP51) and the accumulation of eburicol, which is in full agreement with our hypothesis. Based on our previous finding that inhibition of the cell wall patch formation delays azole-induced fungal death ^7^, our present results support a model where especially the buildup of eburicol is strikingly toxic for *A. fumigatus* (Fig. 7 D). This model is further supported by our finding that lowered expression of Erg6A results in increased azole resistance. Mechanistically, this means that repression of *erg6A* leads to reduced conversion of lanosterol to eburicol, and therefore less CYP51 is needed to diminish accumulating eburicol.

But why did the lowered expression of the Erg9 and Erg1 not result in a similar increase in azole resistance? Instead, we found that it results in a slightly increased azole susceptibility. We believe that this is linked with the overall reduced synthesis of sterols which is expected when the enzymatic turnover by these enzymes is reduced. Because of this, the fungus will not only suffer from the accumulation of eburicol but also from a general depletion of sterols which in itself is incompatible with viability. In contrast, lowered expression of Erg6A would retain *de novo* synthesis of lanosterol, which may fulfill the functionality of the needed sterols to a certain degree, mitigating the deleterious effects of azole-induced eburicol accumulation ^29^.

Notably, in our study we found that Erg6A is essential for the viability of *A. fumigatus*. In contrast, Erg6A homologes of yeasts are not essential for viability ^22,30,31^. Consequently, the sterol C24-methyltransferase could represent a specific antifungal target in fungi that have an ergosterol biosynthesis pathway similar to that of *A. fumigatus*, similar as it was proposed recently ^32^.

A key question that arose in the present study was whether it is the accumulation of eburicol, or of the subsequently formed “toxic diol” which actually is responsible for the detrimental effects of azoles in *A. fumigatus*, including the formation of the cell wall patches. In yeasts, the toxic effect of azoles is clearly attributed to the accumulation of the “toxic diol” as deletion of or mutations in *erg3*, which is required to form it, results in azole resistance ^14–17^. However, the conditional ERG3 triple mutant under repressed conditions still formed cell wall carbohydrate patches after azole exposure, even though it does not produce the “toxic diol”. This demonstrates that neither ERG3 nor the “toxic diol” are required for the azole-induced cell wall patch formation. Furthermore, we show that a mutant lacking ERG3 is significantly more susceptible to azoles compared to the wild type. This indicates that the “toxic diol” which also accumulates in azole-treated *A. fumigatus* as a derivative of the toxic eburicol does not contribute to the toxic effect. In contrast, it even suggests that the conversion of eburicol to 14-methylergosta-8,24(28)-dien-3*β*,6α-diol (“toxic diol”, **16**) contributes to the “detoxification” of eburicol. However, it cannot be excluded that the increased azole susceptibility of the ERG3 triple mutant under repressed conditions is linked to other effects, for example, to a more toxic effect of the eburicol when accumulating in combination with ergosta-7,22,24(28)-trien-3β-ol (**12**), 5-dihydroergosterol, or episterol (**11**). In either case, our results show that the toxicity of azoles in *Aspergillus* has a different mechanism than in yeasts.

Finally, we had previously shown that azole concentrations, which are normally fungicidal for *A. fumigatus*, become primarily fungistatic against *A. fumigatus* if the mitochondrial complex III is disrupted or not functional ^7,28^. This correlates with a strikingly reduced formation of carbohydrate patches at these concentrations. The analysis of the sterol profiles of two mutants which lack mitochondrial complex III functionality, due to to loss of the Rieske protein or cytochrome *c*, revealed that upon azole exposure the accumulating eburicol is much more efficiently converted to the “diol” (14-methylergosta-8,24(28)-dien-3β,6α-diol (**16**)) compared to azole-treated wild type. This supports our model, where the conversion of eburicol into 14-methylergosta-8,24(28)-dien-3β,6α-diol (**16**) helps *A. fumigatus* to survive lower azole concentrations and provides an explanation why *A. fumigatus* mutants with a dysfunctional mitochondrial complex III are more tolerant to azole antifungals. Furthermore, it supports the presumed fungistatic effects attributed to the accumulation of the “toxic diol”, which is regularly observed in azole-treated yeasts. In summary, our results demonstrate that the strong antifungal activity of azoles antifungals on the fungal pathogen *A. fumigatus* relies on the specific accumulation of toxic eburicol which triggers the formation of cell wall carbohydrate patches that contribute to the strong fungicidal activity.

## MATERIAL AND METHODS

### Strains, culture conditions and chemicals

The nonhomologous end joining-deficient strain AfS35, a derivative of *A. fumigatus* D141 ^33,34^, served as wild-type strain for all mutants used and constructed in this work. Conditional mutants and deletion mutants were constructed essentially as described before ^20,34^. In all newly constructed deletion mutants, the respective genes were replaced by double-crossover homologous recombination using a self-excising hygromycin B resistance ^35^. In all newly constructed conditional mutants, a doxycycline-inducible *pkiA*-*tetOn* promoter cassette ^21^ was inserted upstream of the coding sequence of the respective genes. Mitochondria were visualized with a mitochondria-targeted green fluorescent protein (mtGFP) by transforming the respective strains with pCH005, essentially as described before ^21^. All *A. fumigatus* strains used in this study are listed in Table 1. The *Candida* strains used were *C. albicans* ATCC14053 and *C. glabrata* ATCC2950. *Aspergillus* minimal medium (AMM) ^36^ and Sabouraud medium [4% (w/v) D-glucose, 1% (w/v) peptone (#LP0034; Thermo Fisher Scientific; Rockford, IL, US), pH 7.0±0.2] were used in this study. Strains were generally maintained on AMM to obtain new conidia. Solid media were supplemented with 2% (w/v) agar (#214030; BD Bioscience, Heidelberg, Germany). Calcofluor white (Fluorescent brightener 28; #ICNA0215806705) was obtained from VWR International (Radnor, PA, USA). Etest strips were obtained from bioMérieux (voriconazole; #412490; Marcy-l’Étoile, France). Doxycycline-hyclate was obtained from Sigma-Aldrich (#D9891-5G; St. Louis, MO, USA). Voriconazole was obtained from Biorbyt (#orb134756, Cambridge, East of England, United Kingdom). Terbinafin was obtained from Sigma-Aldrich (#T8826, St. Louis, MO, USA ).

**Table 1.** *A. fumigatus* strains relevant for or used in this work.

| Strain or genotype | Relevant genetic modification | Parental strain | Reference |
| --- | --- | --- | --- |
| AfS35 (wt) | *akuA::loxP* | D141 | 33,34 |
| *erg6A<sub>tetOn</sub>* | *(p)erg6A::ptrA-(p)pkiA-tetOn* | AfS35 (wt) | This work |
| *erg6A<sub>tetOn</sub>* + mtGFP | pCH005 | *erg6A<sub>tetOn</sub>* | This work |
| Δ*erg6B* | *erg6B::loxP-hygroR/blaster* | AfS35 (wt) | This work |
| *cyp51A<sub>tetOn</sub>*<br>Δ*cyp51B* | *cyp51B::loxP-hygroR/blaster* | *cyp51A<sub>tetOn</sub>* | <sup>7</sup> |
| *cyp51A<sub>tetOn</sub>* | *(p)cyp51A::ptrA-(p)pkiA-tetOn* | AfS35 (wt) | <sup>7</sup> |
| *cyp51A<sub>tetOn</sub>*<br>Δ*cyp51B* + mtGFP | pCH005 | *cyp51A<sub>tetOn</sub>*<br>Δ*cyp51B* | <sup>7</sup> |
| *erg9<sub>tetOn</sub>* | *(p)erg9::ptrA-(p)pkiA-tetOn* | AfS35 (wt) | This work |
| *erg1<sub>tetOn</sub>* | *(p)erg1::ptrA-(p)pkiA-tetOn* | AfS35 (wt) | This work |
| wt + mtGFP | pCH005 | AfS35 (wt) | <sup>7</sup> |
| Δ*erg3A* | *erg3A::loxP-hygroR/blaster* | AfS35 (wt) | This work |
| Δ*erg3B* | *erg3B::loxP-hygroR/blaster* | AfS35 (wt) | This work |
| Δ*erg3C* | *erg3C::loxP-hygroR/blaster* | AfS35 (wt) | This work |
| *erg3A<sub>tetOn</sub>* | *(p)erg3A::ptrA-(p)pkiA-tetOn* | AfS35 (wt) | This work |
| *erg3A<sub>tetOn</sub>* Δ*erg3B* | *erg3B::loxP-hygroR/blaster* | *erg3A<sub>tetOn</sub>* | This work |
| *erg3A<sub>tetOn</sub>* Δ*erg3C* | *erg3C::loxP-hygroR/blaster* | *erg3A<sub>tetOn</sub>* | This work |
| *erg3C<sub>tetOn</sub>* | *(p)erg3C::ptrA-(p)pkiA-tetOn* | AfS35 (wt) | This work |
| *erg3C<sub>tetOn</sub>* Δ*erg3B* | *erg3B::loxP-hygroR/blaster* | *erg3C<sub>tetOn</sub>* | This work |
| *erg3A<sub>tetOn</sub>* Δ*erg3B*<br>Δ*erg3C* | *erg3C::loxP-hygroR/blaster* | *erg3C<sub>tetOn</sub>*<br>Δ*erg3B* | This work |
| *cycA<sub>tetOn</sub>* | *(p)cycA::ptrA-(p)pkiA-tetOn* | AfS35 (wt) | <sup>7</sup> |
| *rip1<sub>tetOn</sub>* | *(p)rip1::ptrA-(p)pkiA-tetOn* | AfS35 (wt) | <sup>7</sup> |

### Microscopy

Microscopy was performed with a CSU-W1 Spinning Disk Field Scanning Confocal System (Yokogawa; Tokyo, Japan) mounted on a Eclipse Ti2 Inverted Microscope System (Nikon Instruments Inc; Melville, NY, USA) with the exception of the microscopy of terbinafine-treated hyphae, which was performed with a Lionheart FX automated microscope (Agilent BioTek; Santa Clara, CA, USA). For spinning disc confocal microscope, conidia were inoculated in 15 μ-Slide eight-well (#80826) slides (Ibidi; Martinsried, Germany) in 300 µl medium per well. Hyphae that were analyzed with the Lionheart FX automated microscope were inoculated in 96-well plates (#167008; Thermo Fisher Scientific; Rockford, IL, US) in 200 µl medium per well. Samples were generally incubated in humidified chambers to avoid evaporation of medium. Cell wall chitin was stained with calcofluor white. When indicated, samples were fixed with 3.7% formaldehyde in ddH_2_O for 3 min. These samples were then stained with 10 mg ml−1 calcofluor white in ddH_2_O for approximately 1 min, followed by washing the wells three times with ddH2O. For live-cell microscopy and analysis of non-fixed samples, hyphae were stained by supplementing the medium with 3.33 μg ml^−1^ calcofluor white for at least 5 min. The viability of hyphae expressing mitochondria-targeted GFP was analyzed by recording short time-lapse sequences with the spinning disc confocal microscopy, essentially as described before 7. Hyphal compartments with evident mitochondrial dynamics were counted as viable.

### Sterol analysis

Approximately 1 × 10^6^ conidia were inoculated in 50 ml medium per flask and sample. For each condition, three biological replicates of the respective strain were cultivated and analyzed separately. Flasks were incubated in rotary shakers at 37 °C at 180 rounds per minute. To remove doxycycline, the mycelium of each sample was transferred into a Miracloth filter (#475855-1R, Merck Millipore, Brulington, MA, USA) fitted in a funnel and washed three times with medium. The mycelium was then transferred into a new bottle with fresh medium. To harvest mycelium for analysis by gas chromatography-mass spectrometry (GC-MS), mycelium was transferred into a Miracloth filter in a funnel, washed three times with phosphate-buffered saline, frozen in liquid nitrogen, crushed with a mortar and pestle and stored at -80 °C. The thawed wet fungal biomass was subsequently used for sterol extraction as described by Müller *et al.* ^11,37^.

The sterols were analyzed as their corresponding trimethylsilyl (TMS) ethers by GC-MS, essentially as described before ^11^. Overall, 15 different sterols were detected. The sterol TMS ethers were identified by mass spectra and relative retention times (RRT). The base peak of each sterol TMS ether were taken as a quantifier ions for calculating the peak areas for internal standard (IS) cholestane *m/z* 217, RRT 1.00; ergosta-8,22,24(28)-trien-3β-ol *m/z* 253, RRT 1.28; ergosta-5,8,22-trien-3β-ol (lichesterol) *m/z* 363, RRT 1.30; ergosta-5,8,22,24(28)-tetraen-3β-ol *m/z* 361, RRT 1.31; ergosta-5,7,22-trien-3β-ol (ergosterol, **12**) *m/z* 363 RRT 1.33; ergosta-7,22-dien-3β-ol (5-dihydroergosterol) *m/z* 343, RRT 1.36; ergosta-5,7,22,24(28)-tetraen-3β-ol (dehydroergosterol, **13**) *m/z* 361, RRT 1.36; ergosta-8,24(28)-dien-3β-ol (fecosterol, **10**) *m/z* 365, RRT 1.37; ergosta-7,22,24(28)-trien-3β-ol (**12**) *m/z* 343, RRT 1.38; ergosta-5,7-dien-3β-ol *m/z* 365, RRT 1.40; ergosta-7,24(28)-dien-3β-ol (episterol, **9**) *m/z* 343, RRT 1.41; 14-methylergosta-8,24(28)-dien-3β,6α-diol (“toxic diol”, **16**) *m/z* 377, RRT 1.42; 4,4,14-trimethylcholesta-8,24(25)-dien-3β-ol (lanosterol, **1**) *m/z* 393, RRT 1.44; 4-methylergosta-8,24(28)-dien-3β-ol (**5**) *m/z* 379, RRT 1.46; 4,4,14-trimethylergosta-8,24(28)-dien-3β-ol (eburicol, **2**) *m/z* 407, RRT 1.50; 4,4-dimethylergosta-8,24(28)-dien-3β-ol (**4**) *m/z* 408, RRT 1.53. The peak area of each sterol TMS ether was determined, and the percentage of sterol was calculated ^10,38^. Statistical significance was calculated with GraphPad Prism (Version 10.1.1 (Prism 10); Dotomatics, Boston, MA, USA) was calculated with a two-way ANOVA with Tukey’s multiple comparison test for all sterols.

### Use of generative artificial intelligence

DeepL (deepl.com; Cologne, Germany) was used to identify and correct errors in spelling, grammar and punctuation in the manuscript texts.

## ACKNOWLEDGMENTS

This work was in part supported by the German Research Foundation (DFG - WA 3016/4-1), the Irish Health Research Board (SS-2022-016), the Förderprogramm für Forschung und Lehre (FöFoLe) of the Medical Faculty of the LMU München, and the Graduate School of Life Sciences (GSLS) at the University of Würzburg. C.M. thanks Franz Bracher for providing his laboratories and equipment.

---

**Supplementary Fig. 1.**
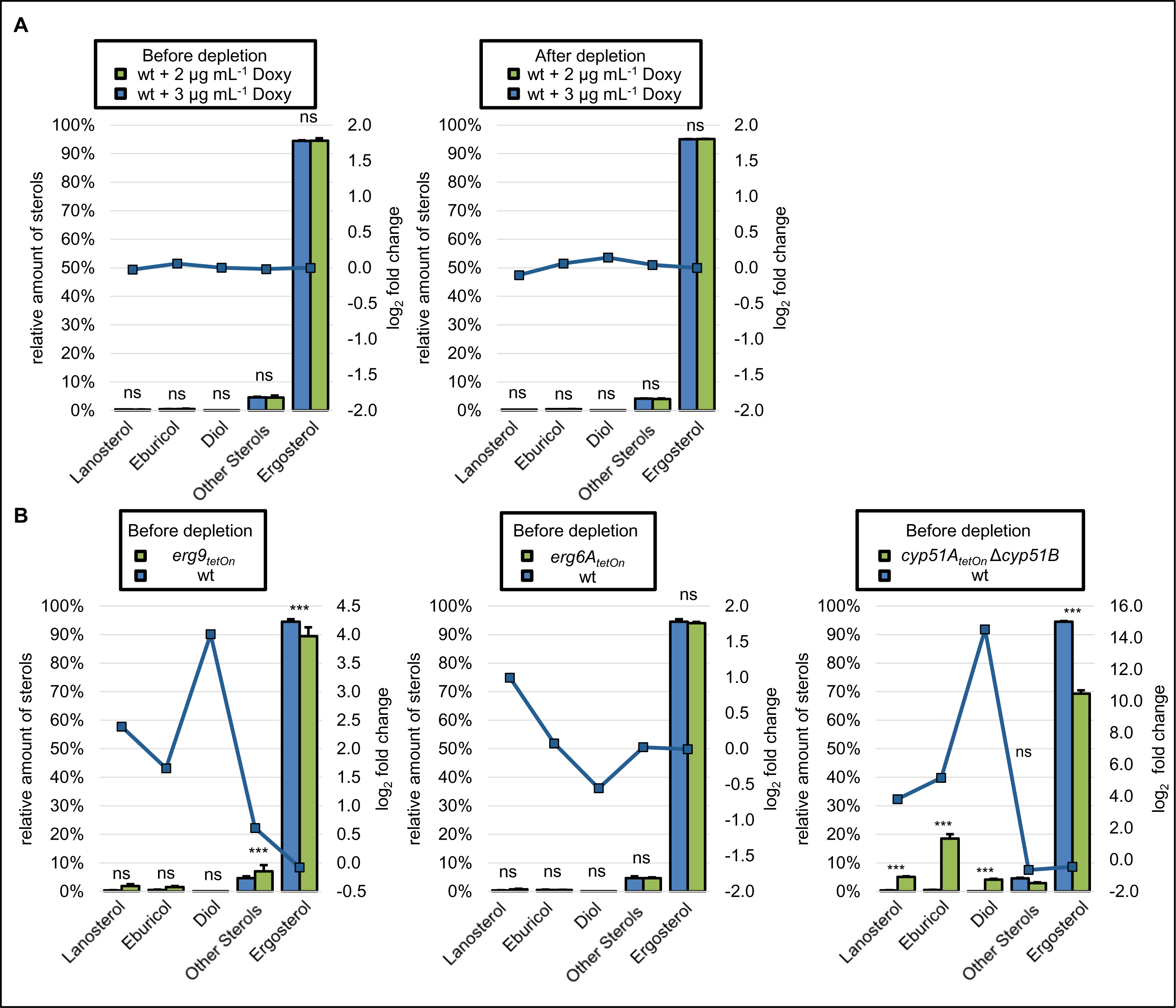
Sterol profiles of squalene synthase-, sterol C24-methyltransferase- and sterol C14-demethylase-depleted *A. fumigatus* hyphae. Further pairwise comparisons of the results of the experiment described and depicted in Fig. 4. The sterol patterns of the harvested mycelia were analyzed by gas chromatography-mass spectrometry (GC-MS). The column graphs show the relative amounts (percentage of total sterol, left y-axis) of the indicated sterols for the indicated strains before and after doxycycline depletion. The data points with the square symbols indicate the log_2_-fold change (right y-axis) in the amount of the respective sterol of the pairwise comparison of the conditions shown in the individual graphs. The log_2_-fold change data points were connected by lines for better visual illustration of the changes in the profiles. The column bar data points represent means of three replicates per condition, the error bars indicate standard deviations. Statistical significance (* p<0.05; ** p<0.01; *** p<0.001) was calculated with a two-way ANOVA with Tukey’s multiple comparison test.

